# A mobile ESX type VII secretion system enhances intracellular persistence in globally distributed *Mycobacterium abscessus*

**DOI:** 10.64898/2026.01.26.701661

**Authors:** Kia C. Ferrell, Andrew H. Buultjens, Sherridan Warner, Sibel Alca, Andrea Bustamante, Eby M. Sim, Elena Martinez, Vitali Sintchenko, Claudio Counoupas, Timothy P. Stinear, James A. Triccas

## Abstract

*Mycobacterium abscessus* are non-tuberculous mycobacteria that are widespread in the environment and of increasing global clinical significance. Accumulating evidence shows that *M. abscessus* has emerged as an important pathogen, driven by highly drug-resistant lineages, enhanced transmissibility and the acquisition of specific virulence factors. In this study, we describe a previously uncharacterised ESX secretion system encoded on a 123-kbp plasmid identified in a clinical isolate of *M. abscessus*. This ESX system, termed ESX-pMA07, is distinct from ESX systems previously reported in *M. abscessus* in both sequence composition and locus organisation, characterised by a unique arrangement of core ESX components and low sequence identity to ESX-3, ESX-4 and plasmid-borne ESX-P systems. ESX-pMA07 was detected in geographically diverse clinical isolates but was restricted to particular genotypes within the global *M. abscessus* phylogeny. Transcriptional profiling revealed expression of ESX-pMA07 components in artificial cystic fibrosis media and during intracellular growth in macrophage cell lines. Using CRISPR interference, we show that inducible silencing of *eccC*, encoding the ATPase component of ESX-pMA07, significantly reduced intracellular survival of *M. abscessus* within macrophages. To our knowledge, this is the first characterisation of a functional, plasmid-borne ESX secretion system in *M. abscessus*, demonstrating that mobile genetic elements contribute to the pathogen’s intracellular persistence and may influence its evolving virulence.

**Author Summary:** *Mycobacterium abscessus* is a rapidly emerging, highly drug-resistant bacterium that causes chronic infections, particularly in people with underlying lung disease such as cystic fibrosis. The factors that enable certain *M. abscessus* strains to persist inside host cells are not fully understood. In this study, we identified a previously unrecognised type VII secretion system (ESX) encoded on a large plasmid in a clinical isolate of *M. abscessus*. This plasmid-borne ESX system, which we termed ESX-pMA07, is genetically distinct from the ESX systems normally found on the chromosome and was detected in geographically diverse clinical isolates, but restricted to specific lineages within the global *M. abscessus* population. We show that ESX-pMA07 genes are expressed under conditions relevant to lung infection and during intracellular growth in macrophages. Using inducible CRISPR interference to silence the ESX ATPase gene *eccC*, we demonstrate that ESX-pMA07 contributes to intracellular survival of *M. abscessus* in macrophages. These findings reveal that mobile genetic elements can encode functional secretion systems that enhance intracellular persistence, providing a mechanism for the emergence and spread of virulence traits in this important pathogen.

## Introduction

*Mycobacterium abscessus* (also known as *Mycobacteroides abscessus*) is a multi-species complex within the *Mycobacterium*, a genus that harbours both ubiquitous, environmental bacteria and major human pathogens [1]. Species within the *M. abscessus* complex are increasingly found in urban environments such as drinking water distribution systems and shower heads [2]. This encroachment of non-tuberculous mycobacteria (NTM) such as *M. abscessus* into human environments has increased the risk for *M. abscessus* to cause opportunistic infections (Falkinham, 2009).

NTM primarily cause skin and soft tissue infections, but they can also cause chronic pulmonary infections, known as NTM pulmonary disease (NTM PD), in susceptible people [3]. People with reduced pulmonary immune responses such as cystic fibrosis (CF) or chronic obstructive pulmonary disease (COPD) are particularly susceptible to NTM PD. Globally, incidence rates of NTM infections including those caused by *M. abscessus* are increasing [4–6]. Unfortunately, the host and pathogen factors leading to chronic infections caused by NTM - as opposed to transient NTM infection and pathogen clearance - remain unclear [4,7]. Infection with NTM is associated with continued disease progression and decline in lung function [8]. For those affected by NTM PD, infection is extremely difficult to treat due to extensive resistance of *M. abscessus* to antibiotics and disinfectant agents commonly used in nosocomial settings [9,10]. There are no vaccines to prevent or treat *M. abscessus* infections and resistance to antibiotics is common [10]. Thus, there is an urgent need for preventative and therapeutic treatments for this pathogen.

*M. abscessus* has diverse virulence attributes including immunomodulatory factors, some of which are also shared with *Mycobacterium tuberculosi*s, the causative agent of human tuberculosis [11,12]. Unlike *M. tuberculosis*, there is evidence suggesting *M. abscessus* has also acquired virulence factors from bacterial species of distant genera such as *Pseudomonas aeruginosa*, that are also responsible for chronic bacterial infections in susceptible individuals, such as CF patients [13,14]. The genomic plasticity of *M. abscessus* is concerning as infection rates rise [4,15,16]. The emergence of dominant circulating *M. abscessus* clones that have increased virulence in animal infection models and show evidence of adaptation to the physiological conditions of the human lung are of particular concern [17,18]. Furthermore, the widespread presence of mobile genetic elements such as plasmids and prophages within *M. abscessus* suggests there is a high potential transmission of pathogenic traits within and among the complex [19]. The expansion of the CF patient population over the last century has also increased the reservoir of highly susceptible potential hosts for *M. abscessus*, and this has perhaps exacerbated the rate of pathogen evolution [20].

In this study we utilised *in silico* analysis to screen *M. abscessus* clinical isolates for novel virulence traits. Comparative genome analysis of *M. abscessus* clinical isolate MA07 identified a novel, plasmid borne Type VII secretion system (T7SS; ESX secretion system) termed ESX-pMA07 that was present in geographically diverse clinical isolates of *M. abscessus*. Components of this ESX secretion system were expressed during *in vitro* growth in laboratory conditions, including culture in artificial CF media and during intracellular infection of macrophages. Using an inducible CRISPR silencing approach, reversible suppression of the putative ATPase *eccC* within the ESX-pMA07 secretion system resulted in decreased intracellular survival of *M. abscessus* in macrophages. This study is the first description of a plasmid borne ESX system in *M. abscessus* that contributes to intracellular replication and is thus likely contributing to the virulence of this pathogen.

## Results

### MA07 clinical isolate has a plasmid-borne ESX secretion system

A screen of *M. abscessus* clinical isolates identified a strain, termed MA07, that displayed heightened replication in a macrophage model of *M. abscessus* infection (Figure S1). To find bacterial factors contributing to this enhanced persistence, we determined the complete genome sequence of clinical isolate MA07 using both short and long-read sequencing and compared it to other mycobacterial genomes. MA07 has a 5,112,298 bp genome comprising a single circular chromosome and a 123,141 bp plasmid. Virulence factor (VF) analysis comparing MA07 predicted protein sequences with VFs in *M. abscessus* ATCC 19977 (GenBank accession: CU458896.1) and *M. tuberculosis* H37Rv (GenBank accession: AL123456.3) identified 149 putative VFs in MA07, 135 in ATCC 19977 and 253 in H37Rv (Figure 1). Of particular interest, MA07 contained proteins with similarity to the mycobacterial T7SS ESX-3 and ESX-5. These genes mapped to a single locus, present on the MA07 plasmid that was named pMA07. This plasmid contained 151 predicted coding sequences, including the ESX secretion system components identified and a putative origin of replication (Figure 2A). *In silico* comparisons showed limited sequence similarity between pMA07 and previously described ESX-containing plasmids (Figure 2B, Table S1).

**Figure 1.**
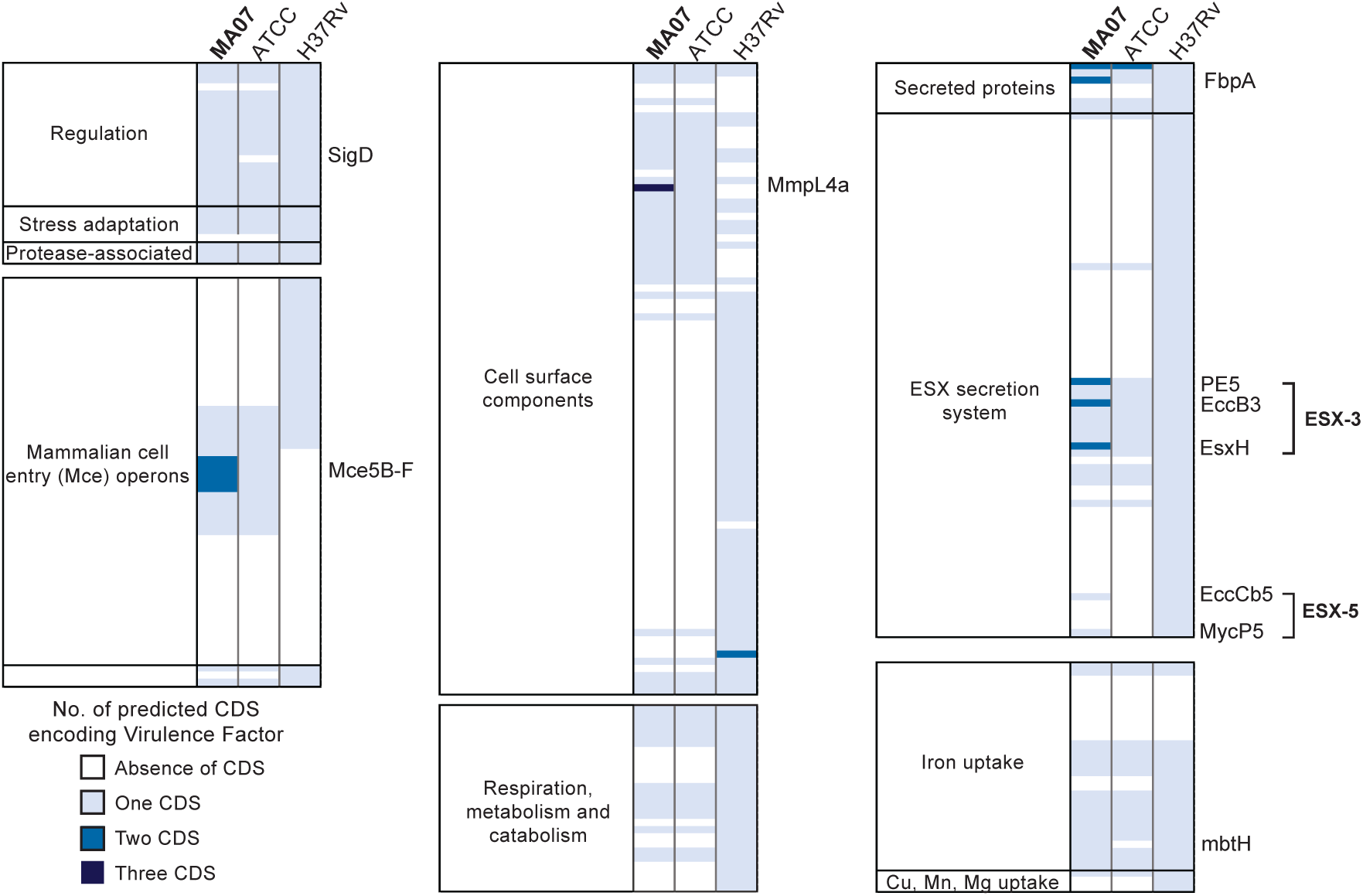
Novel potential virulence factors within clinical isolate *M. abscessus* MA07. Identification of novel putative virulence factors within the MA07 genome identified by VF analyser. Degree of shading corresponds to the number of coding sequences (CDS) present within *M. abscessus* MA07 (MA07), reference strain *M. abscessus* ATCC19977 (ATCC) and *M. tuberculosis* H37Rv (H37Rv). Virulence factors present in MA07 that are absent in the reference strain are annotated with their abbreviated name.

**Figure 2.**
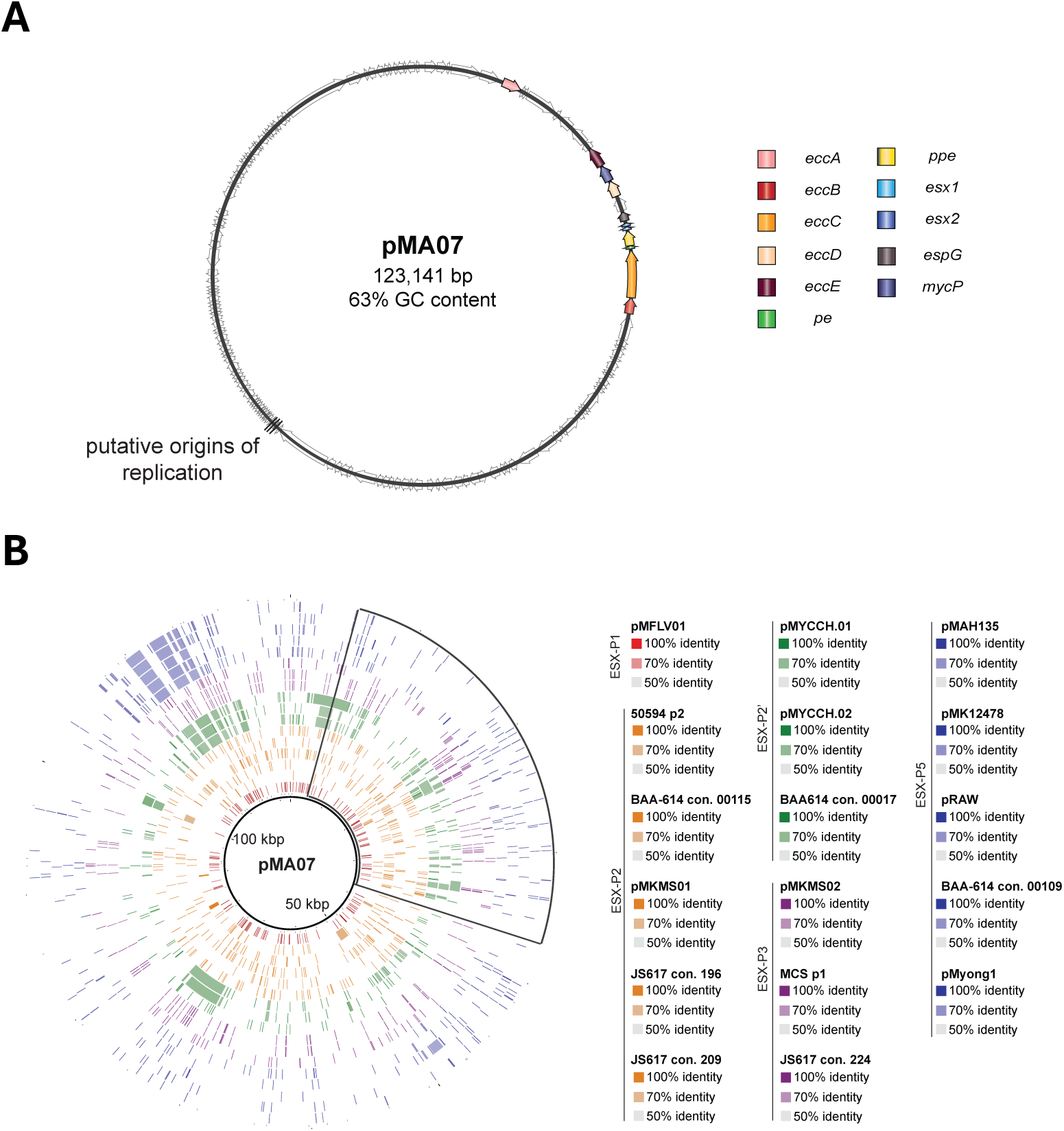
*M. abscessus* MA07 contains a plasmid borne ESX secretion system. **(A)** Schematic showing pMA07 and relative positions of ESX locus and origin of replication. pMA07 was identified by Oxford Nanopore long read sequencing, assembly and annotation of MA07 DNA. Position of ESX locus on pMA07 was visualised using SnapGene, and other coding regions are represented by grey arrows. Putative origin of replication identified by OriFinder. **(B)** Similarity of pMA07 sequence to other plasmids or contigs containing ESX secretion systems, determined by BLAST Ring Image Generator (BRIG) analysis. Plasmids are listed in order of display (outwards) and colours correspond to different classes of plasmid borne ESX secretion systems as described by Dumas et al. 2016. Outlined region corresponds to the location of the ESX secretion system within pMA07, and central ruler indicates plasmid position in kilobase pairs (kbp).

Detailed BLASTp analysis of the annotated plasmid suggested the pMA07 ESX locus contained the components of a functional T7SS. Five ESX conserved component proteins (EccA-E), three secretion targets (PE, PPE and an WXG100 protein we termed Esx2), one ESX specific protein (EspG) and a serine protease mycosin MycP were identified (Figure 3). In addition, SWISS-MODEL analysis showed that MA07_4597 encoded a hypothetical protein with a tertiary structure similar to the ESX-1 secretion component EsxB, which we named Esx1. All identified predicted proteins had low (less than 50%) amino acid identity to orthologous proteins in reference strain *M. abscessus* ATCC19977 (Table 1), and had a genomic organisation of ESX components that was distinct to other previously described ESX-3, ESX-4 and ESX-P systems (Figure 3).

**Figure 3.**
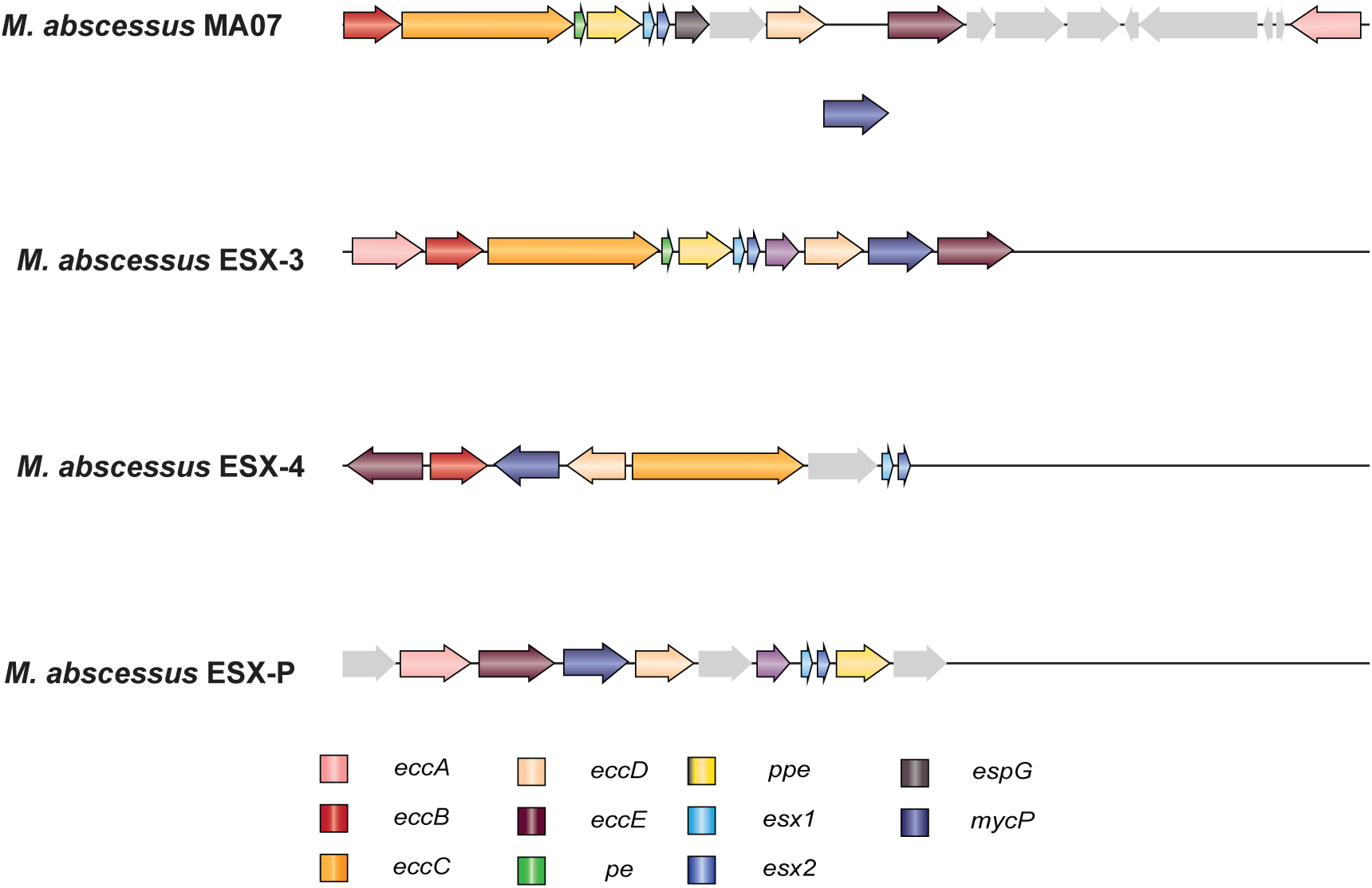
*M. abscessus* pMA07 contains a novel ESX secretion system. Gene map showing the layout of *M. abscessus* pMA07 secretion system, in comparison with *M. abscessus* ESX-3, ESX-4 and plasmid borne ESX-P. Genes of unknown function are represented in grey; length of coding sequences shown is representative and is not to scale. Schematics for ESX-3, ESX-4 and ESX-P were adapted from Johansen et al, 2021.

**Table 1.** ESX secretion system components identified by BLASTp within *M. abscessus* MA07 region of difference

| **Protein name** | **Gene identity** | **BLASTp hit** | **% sequence coverage to *M. abscessus* ATCC19977** | **% sequence identity to *M. abscessus* ATCC 19977** |
| --- | --- | --- | --- | --- |
| **EccB** | MA07_4593 | type VII secretion protein EccB<br>[*Mycobacteroides abscessus*] | 95 | 32.98 |
| **EccC** | MA07_4594 | type VII secretion protein<br>EccCa [*Mycobacteroides abscessus*] | 90 | 30.1 |
| **PE** | MA07_4595 | PE domain-containing protein<br>[*Mycobacteroides abscessus*] | No significant similarity found |  |
| **PPE** | MA07_4596 | PPE domain-containing protein<br>[*Mycobacteroides abscessus*] | 7 | 48.48 |
| **Esx1** | MA07_4597 | hypothetical protein<br>[*Mycobacteroides abscessus*] | No significant similarity found |  |
| **Esx2** | MA07_4598 | WXG100 family type VII<br>secretion target<br>[*Mycobacteroides abscessus*] | No significant similarity found |  |
| **EspG** | MA07_4599 | ESX secretion-associated<br>protein EspG [*Mycobacteroides abscessus*] | 43 | 33.87 |
| **EccD** | MA07_4601 | type VII secretion integral<br>membrane protein EccD<br>[*Mycobacteroides abscessus*] | No significant similarity found |  |

### Distribution of pMA07 and ESX-pMA07 are restricted among a global *M. abscessus* population

Plasmids play a key role in the dissemination and evolution of ESX systems in mycobacteria, acting as vehicles of horizontal transfer of ESX associated genes [21]. We therefore reasoned that the pMA07 plasmid may be present across the *M. abscessus* clade [21,22]. To examine the distribution of pMA07 and its associated T7SS, we inferred a global phylogeny using 1,160 publicly available *M. abscessus* genome datasets (Figure 4). For each isolate, we assessed read-mapping coverage across all 151 pMA07 genes and defined “presence” of pMA07 plasmid sequences as ≥50% gene presence. Using this criterion, we identified 10 additional strains of *M. abscessus* that harbour substantial portions of pMA07, including the complete ESX-pMA07 locus (Figure 4, red shading in outer ring; Table S2). Nine of these 11 isolates clustered within the dominant circulating clone DCC1, a lineage associated with increased virulence and global spread [17,23]. Further analysis revealed that strains containing a high proportion of pMA07 sequences were clinical isolates, most frequently respiratory isolates (7/10), and from geographically diverse origins (Table 2). Protein homology screening for all components of the pMA07 ESX system showed that each strain identified contained all components of the ESX-pMA07 system, with a high degree of sequence similarity and identity (Figure 5).

**Figure 4.**
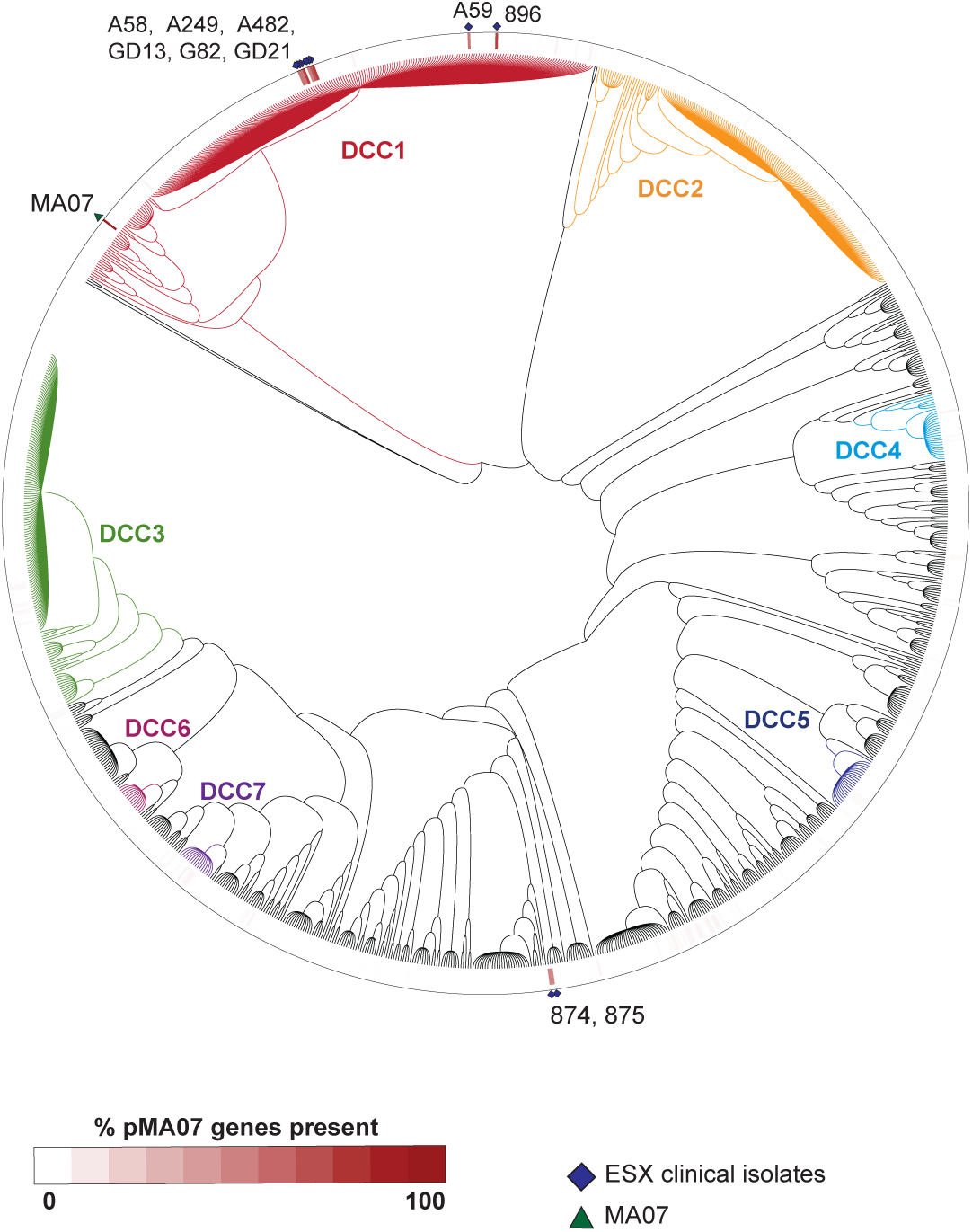
Distribution of pMA07 across *M. abscessus* global phylogeny. Maximum likelihood tree of 1,160 global isolates of *M. abscessus* mapped against pMA07. Tree was inferred from a core-SNP alignment using FastTree and visualised using iTOL. Coloured branches denote identification of seven dominant circulating clones (DCCs) of *M. abscessus*, and shading in outer circular heat map represents the percentage of pMA07 plasmid genes present in each isolate within the phylogeny. Positions of MA07 (green triangle) and clinical isolates containing the ESX system (blue diamonds) are shown in the outer ring of the phylogeny.

**Figure 5.**
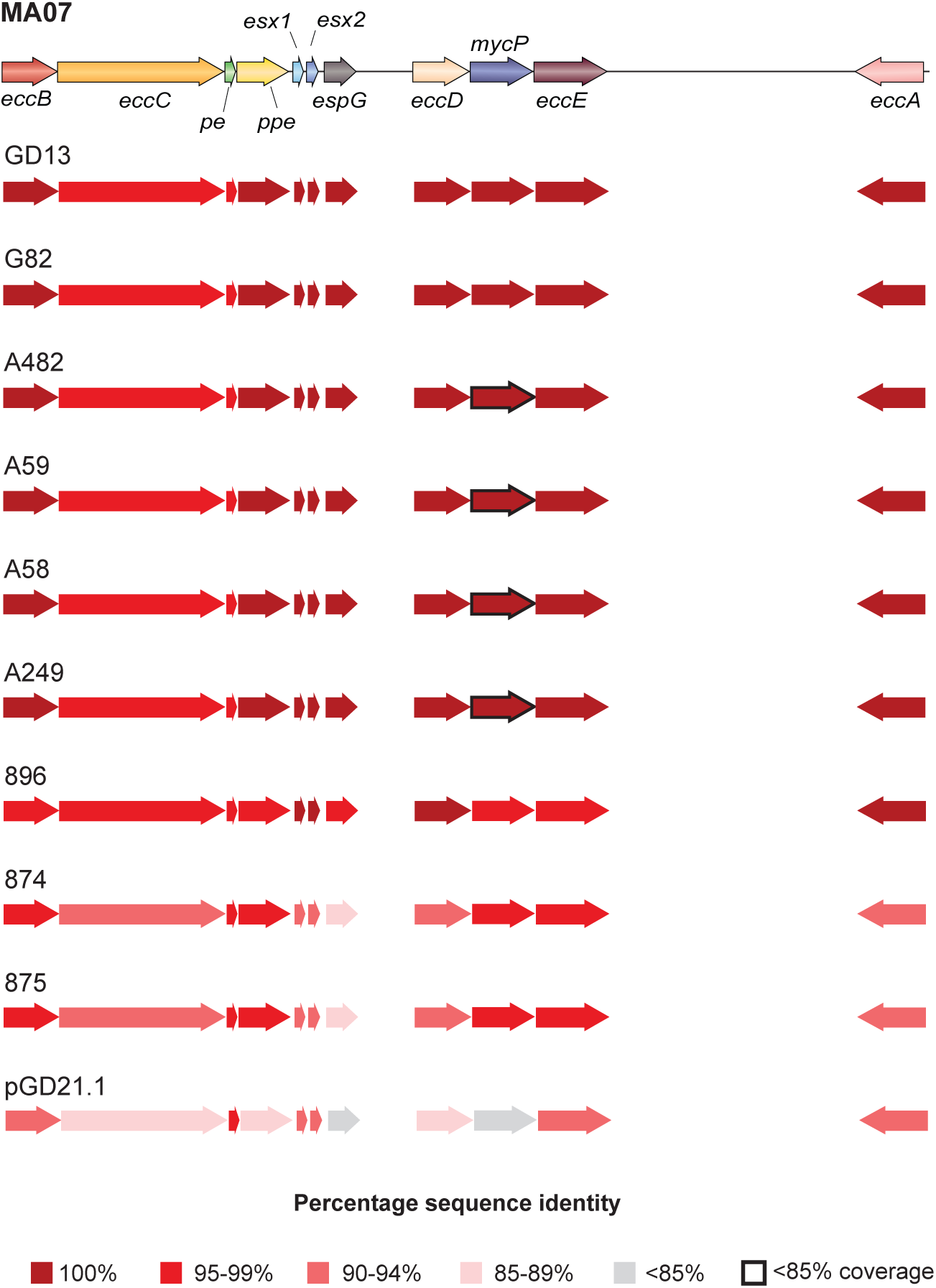
Sequence similarity of *M. abscessus* secretion system among globally diverse clinical isolates. Locus map representing the sequence similarity of different clinical isolates (described in Table 2) to MA07. Similarity was determined by BLASTp analysis; Degree of shading is representative of percentage protein sequence identity (rounded to the nearest whole number); sequences with <85% sequence coverage are represented with a black border.

**Table 2.** Strain details of *M. abscessus* isolates containing ESX secretion system proteins

| Strain name | Clade | Isolate source | Isolate geographical location | Disease manifestation | Identifier (BioSample ID) |
| --- | --- | --- | --- | --- | --- |
| **896** | *M. abscessus*<br>subsp. *abscessus* | Respiratory system (CF) | Ireland | Diseased, *M. abscessus* pulmonary | SAMEA2259701 |
| **G82** | Not specified | Sputum | China | *M. abscessus* lung disease | SAMN09917584 |
| **GD13** | *M. abscessus*<br>subsp. *abscessus* | Clinical patient | Pennsylvania, USA | NTM infection | SAMN16839142 |
| **A59** | *M. abscessus*<br>subsp. *abscessus* | Sputum (non-CF) | Shanghai, China | *M. abscessus* lung disease | SAMN07501872 |
| **A482** | *M. abscessus*<br>subsp. *abscessus* | Lavage fluid (Non-CF) | Shanghai, China | *M. abscessus* lung disease | SAMN07501889 |
| **A58** | *M. abscessus*<br>subsp. *abscessus* | Sputum (Non-CF) | Shanghai, China | *M. abscessus* lung disease | SAMN07501871 |
| **A249** | *M. abscessus*<br>subsp. *abscessus* | Sputum (Non-CF) | Shanghai, China | *M. abscessus* lung disease | SAMN07501881 |
| **875** | *M. abscessus*<br>subsp. *bolletii* | Respiratory system | Australia | Lung disease | SAMEA2259680 |
| **874** | *M. abscessus*<br>subsp. *bolletii* | Respiratory system | Australia | Lung disease | SAMEA2259679 |
| **GD21 (pGD21.1)** | *M. abscessus*<br>subsp. *abscessus* | Clinical patient (Non-CF) | Louisiana, USA | NTM infection | SAMN16828757 |

To determine if the ESX secretion system in pMA07 was present in other bacteria, the sequence of the putative EccD protein (MA07_4601) was used to query all non-redundant proteins within the NCBI BLAST database. Only the 10 *M. abscessus* strains previously identified in the global phylogeny contained a region with 100% coverage and greater than 85% sequence identity to the EccD protein from MA07 (Table S3). Therefore, the ESX-pMA07 secretion system is confined to a limited group of *M. abscessus* isolates identified in our phylogenetic screen, with no detectable homologues in other bacterial genomes.

### pMA07 ESX secretion system components are expressed during *in vitro* culture

We further sought to elucidate the role of ESX-pMA07 by analysis of secretion components during *in vitro* growth of MA07. Expression of *M. abscessus* pMA07 ESX components was measured with sequence-specific primers by qRT-PCR. The specificity of primers used was validated using genomic DNA from MA07 and reference strain CIP104536T, which does not contain this ESX secretion system (Figure 6A). Target sequences were only detected using MA07 template DNA, further validating their specificity to ESX-pMA07. Next, MA07 was cultured *in vitro* in complete 7H9 mycobacterial media and in Sauton’s modified minimal media to determine whether these components were differentially expressed based on their growth environment (Figure 6B). All components, excluding *eccE*, could be detected during *in vitro* growth, and the relative expression of predicted secretion targets PE, PPE, *esx1* and *esx2* was the highest of the different genes. The expression of selected ESX-secreted components was also examined in artificial cystic fibrosis media, which is reflective of the nutrient composition of CF sputum in the lung [24]. High expression of all tested genes was observed in this media, with particularly strong induction of *esx1* and *esx2* compared with both c7H9 and Sauton’s media (Figure 6C).

**Figure 6.**
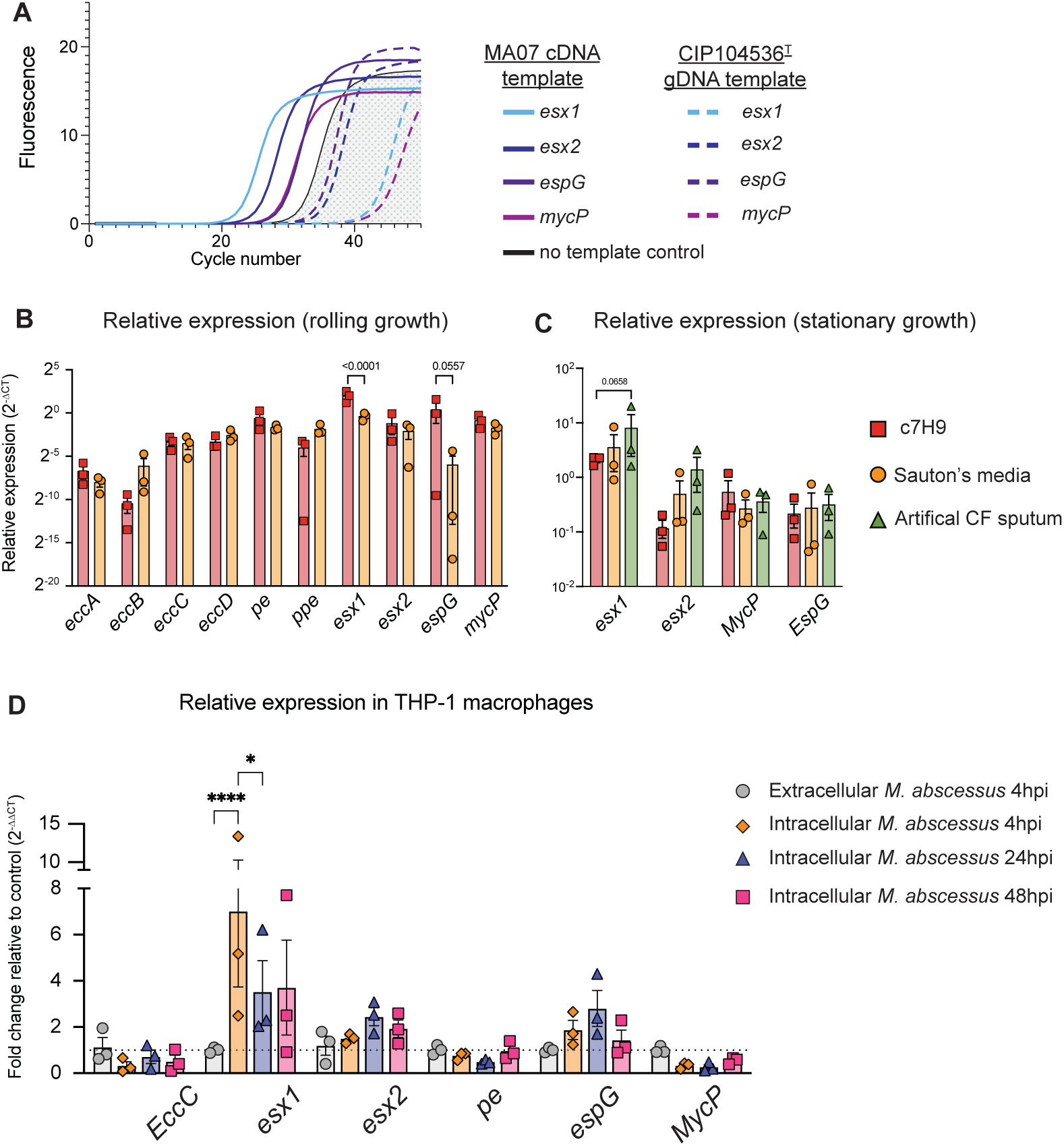
Expression of pMA07 ESX secretion components vary during culture conditions and in macrophages. qRT-PCR amplification curves of ESX components using MA07 cDNA (solid lines) or *M.abscessus* CIP104536T genomic DNA (gDNA; dashed lines) as the template **(A).** Relative expression (2-ΔCT) of ESX secretion components in MA07 grown in complete 7H9 (c7H9) or sauton’s modified media **(B).** Relative expression of ESX components by MA07 during stationary growth in c7H9, Sauton’s modified media or Artificial Cystic Fibrosis (CF) media **(C).** Expression of ESX components in THP-1 macrophages infected with MA07 at 4, 24 and 48hpi, relative to extracellular *M. abscessus* collected at 4hpi **(D).** For all qPCR data, expression is normalised to reference gene sigA and each point represents a biological replicate within one experiment. Graphs are displayed with mean relative expression±SEM. Significance determined by 2-way ANOVA with Šidák’s multiple comparison test (*p<0.05;****p<0.0001).

### *M. abscessus* ESX components alter *M. abscessus* intramacrophage growth

The presence of ESX-pMA07 components among isolates clustering within the virulent DCC1 lineage in the global *M. abscessus* phylogeny led us to hypothesise that this ESX system contributes to *M. abscessus* virulence. To investigate this, we examined the expression of pMA07 ESX components by *M. abscessus* after internalization by THP-1 macrophages, using qRT-PCR (Figure 6D). Of the 6 genes examined, there was a significant increase in expression of *esx1* at 4 hours post infection compared to extracellular *M. abscessus*. There was also a trend towards increased intracellular expression of *esx2* and *espG* at 24 and 48h respectively, but this did not reach statistical significance. Taken together, these results indicate that secretion components are differentially expressed in different growth conditions and during the intracellular stage of *M. abscessus* growth, suggesting a potential role of ESX-pMA07 in *M. abscessus* intracellular survival.

To further define the role of ESX-pMA07 in virulence, an inducible CRISPR knockdown approach was applied to MA07 to silence components of the ESX secretion system (Figure 7A). As the EccC component is critical for substrate transport through the mycomembrane in other mycobacterial T7SS [25,26], this component was targeted for silencing. Guide RNAs were designed for a putative promotor region of the ESX cassette, and multiple locations within the *eccC* locus (figure 7B). sgRNA-EccC6 displayed a reduced normalized expression of all ESX components in the presence of anhydrotetracycline (ATc) during stationary but not rolling growth (Figure S2A-B). One of these knockdowns, EccC6, displayed reduced intracellular survival in macrophages, despite no change in extracellular survival in the presence of ATc (Figure S2C-D). To further clarify the potential role of the EccC protein in the virulence of MA07, intracellular survival of MA07-siRNA EccC6 and empty vector control was monitored in THP-1 macrophages up to 24hpi. Uptake rates of the empty vector control containing no siRNA (EV) and EccC6 by THP-1 macrophages were similar at 4 hours post infection in the presence of ATc (Figure 7C). However, there was a significant reduction in intracellular survival of *M. abscessus*-siRNA EccC6 in THP-1 macrophages at 24 hours (Figure 7D). Taken together, these results suggest a role of EccC, and by extension the ESX-pMA07 system, in intracellular survival of *M. abscessus* MA07.

**Figure 7.**
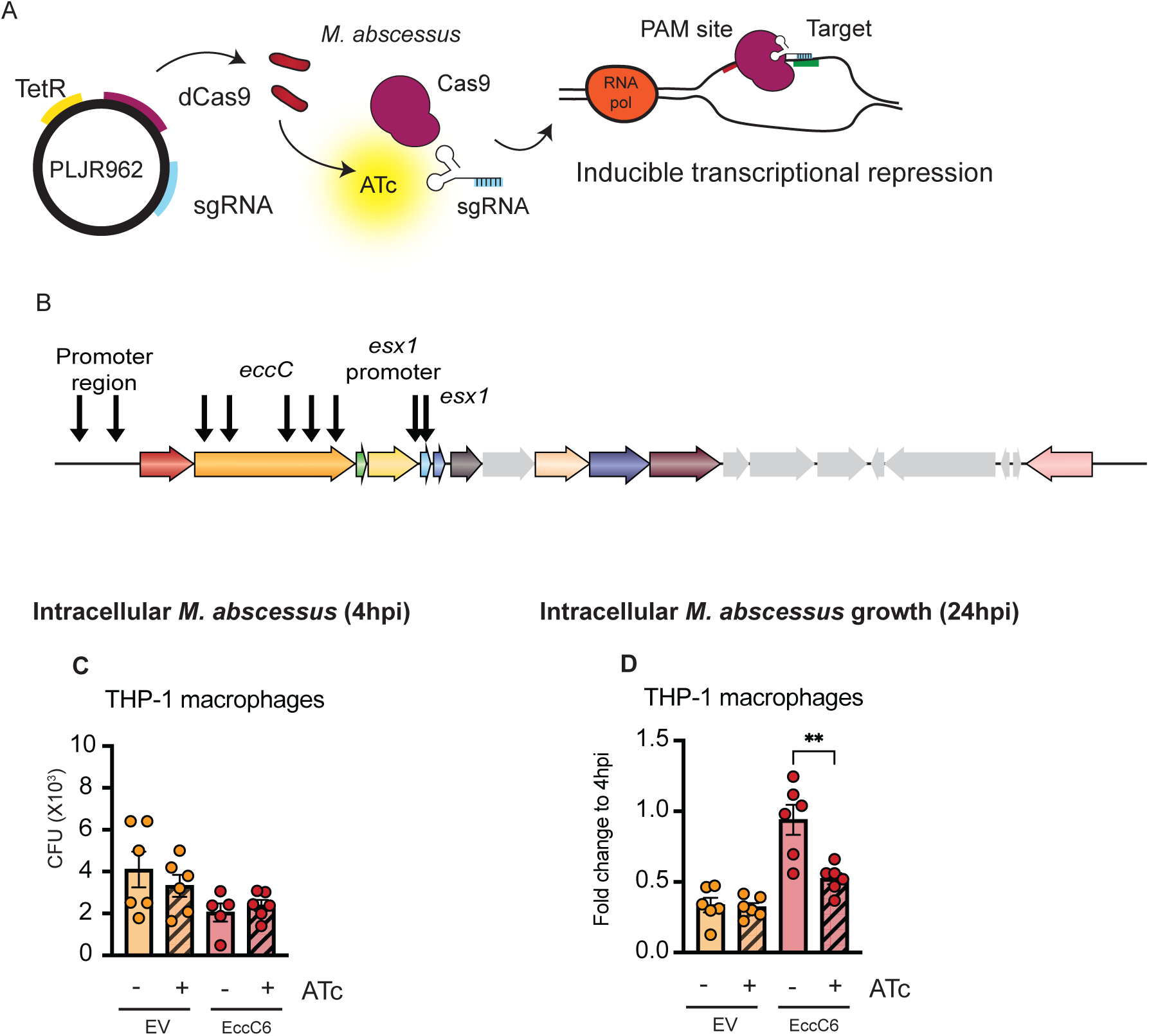
CRISPR inducible silencing of pMA07-EccC reduces *M. abscessus* intramacrophage survival. Schematic detailing the mechanism of inducible transcriptional repression in *M. abscessus* MA07, adapted from Rock *et al.* 2021 Integrative plasmid PLJR962 containing endonuclease deficient Cas9 (dCas9) and guide RNA specific to target gene (sgRNA) under the control of tetracycline repressor (TetR) was transformed by electroporation into MA07. In the presence of anhydrotetracycline (ATc), the CRISPRi complex is formed which binds to the protospacer adjacent motif (PAM) site and target gene, resulting in transcriptional repression **(A).** Location of sgRNA target sequences on ESX locus selected for CRISPRi based transcriptional repression **(B)**. Intracellular bacterial burden of *M. abscessus* MA07-EccC and Empty Vector (EV) transformants at 4 hours post infection **(C)** in THP-1 macrophages in the presence of ATc 500ng/mL; M. abscessus fold change in CFU in THP-1 macrophages **(D)** at 24hpi.

## Discussion

Rising incidence rates of global NTM infections, coupled with the increasing pathogenicity of *M. abscessus,* highlight the urgent need for a deeper understanding of *M. abscessus* host-pathogen interactions. The recent characterisation of *M. abscessus* DCCs underscores a concerning trajectory of *M. abscessus* towards becoming a true human pathogen, accompanied by diverse virulence factors that promote immune evasion and intracellular survival [12,17,27]. The clear link between the acquisition of virulence traits and global spread of *M. abscessus* emphasises the importance of ongoing screening for emergence of new mechanisms of pathogenicity in this species.

Here we describe a novel genetic element within a clinical isolate of *M. abscessus* that enhance virulence. This ESX secretion system is distinct in both sequence and locus organisation from those previously described in *M. abscessus* and other species of mycobacteria [11]. The ESX secretion systems within *M. abscessus*, ESX-3 and ESX-4, have recently been described as playing a significant role in *M. abscessus* virulence by blocking phagosomal maturation and enabling escape to the cytosol [25]. This is consistent with the well-established role of ESX systems as key mediators of mycobacterial virulence [26]. The identification of a functional ESX system on a mobile genetic element therefore has substantial implications, revealing a previously unrecognised mechanism for distributing virulence factors across the *M. abscessus* population.

We identified genomically diverse *M. abscessus* isolates within the global phylogeny that contain the ESX-pMA07 locus. Most of these isolates clustered within DCC1, a genotype associated with chronic infection, heightened survival in macrophages and extensive immunopathology in immune compromised mice [18]. This study found only ten additional isolates across the global phylogeny with a high number of genes corresponding to pMA07 (≥50% gene presence), suggesting relatively limited dissemination of pMA07 within the *M. abscessus* clade. However, detection of plasmid DNA is influenced by the extraction protocol, and commonly used methods such as salting out can lead to loss of plasmids leading to an underestimation of their true prevalence [28,29]. For isolates identified through short-read sequencing, we were unable to determine whether ESX-pMA07 is maintained as a plasmid or has integrated into the chromosome. Thus, mobile genomic elements such as pMA07, which are challenging to identify by short-read sequencing, could have a broader contribution to *M. abscessus* pathogenicity than previously described.

Where bacterial species such as *M. abscessus* harbour plasmids, there is an associated fitness cost that accompanies their retention within the cell. However, it is assumed that the plasmid provides beneficial genes providing some advantage to the host, therefore facilitating its persistence within the population and countering the additional metabolic cost of plasmid maintenance [30,31]. This is particularly significant for large plasmids such as pMA07, which are metabolically costly to maintain in the absence of selective pressure [32], further suggesting the importance of pMA07 to the isolate. While pMA07 likely provides benefit in the circumstance of intracellular growth and virulence, we cannot exclude additional functions. Interestingly, the fitness cost associated with heightened virulence imposes an apparent limitation on *M. abscessus* transmissibility, including decreased survival on fomites [17]. Should the only survival advantage conferred by carrying pMA07 relate to enhanced virulence, this may restrict the continued spread of this plasmid within *M. abscessus*.

The use of CRISPR interference for mycobacteria [33] has been widely used to characterise the function of unknown genes in *M. abscessus* [20,34,35]. The use of CRISPRi based knockdown in this study was selected over homologous recombination approaches [such as used by [36]], which cannot be used to delete genes for multiple copy plasmids. However, we observed a phenotypic change in knockdown *M. abscessus* (i.e. reduced intracellular survival) alongside reduced expression of ESX components during stationary growth, but not during rolling growth (Figure S2). Although knockdown efficiency differed between growth conditions, the intracellular phenotype strongly supports a functional requirement for EccC and reinforces the contribution of ESX-pMA07 to macrophage survival.

The results presented here provide important evidence of the increasing pathogenicity and evolution of *M. abscessus*. The presence of plasmid pMA07 within diverse isolates clustered in the DCC clade is suggestive of the importance of mobile genetic elements in *M. abscessus* virulence. Increased expression of these ESX components within an in vitro setting that reflects the pulmonary environment, together with reduced intracellular survival of this pathogen in murine macrophages following *eccC* silencing, indicates that this novel ESX secretion system comprises a key mechanism of *M. abscessus* virulence that enhances survival within the host. The potential transmissibility of this system on pMA07 highlights a previously unrecognised pathway by which *M. abscessus* may undergo rapid evolutionary gains in pathogenicity.

## Materials and methods

### *M. abscessus* culture conditions

*M. abscessus* isolates were grown in Middlebrook 7H9 media (Beckton Dickinson) supplemented with 10% ADC supplement, 0.2% glycerol and 0.02% tyloxapol while rolling at 37°C. The MA07 isolate was selected from a selection of clinical isolates that displayed heightened persistence in an infection model of *M. abscessus* infection (Supplementary figure 1). Modified Sauton’s minimal media was prepared with (0.4% (w/v) L-Asparagine, 0.1% (w/v) magnesium sulfate heptahydrate, 0.07% (w/v) potassium phosphate trihydrate, 0.2% (w/v) citric acid monohydrate, 0.0005% (w/v) ammonium iron (III) citrate, 0.5% (w/v) sodium pyruvate, 0.0001% (w/v) zinc sulfate heptahydrate, 0.005% (v/v) glycerol and 0.5% (v/v) glucose (Sigma) in Triple Distilled Water (TDW), pH 7.2-7.4. Artificial cystic fibrosis media was prepared as previously described, with some modifications [24]. Briefly, 1% (w/v) porcine mucin, 0.00014% salmon sperm eDNA, 10% BSA, 0.5% egg yolk emulsion, 5% (w/v) casamino acids, 0.003% (w/v) casamino acids, 0.22% (w/v) calcium chloride, 5% (w/v) sodium chloride and 2.2% (w/v) potassium chloride were dissolved overnight while stirring at 4°C before sterile filtration and storage at 4°C. Colony forming units (CFU) were determined by enumeration on LB agar, incubated for 5-7 days at 37°C and 5% CO_2_. Frozen stocks of MA07 were prepared in 7H9 with a final concentration of 25% glycerol and stored at -80°C.

### Virulence factor identification

Putative virulence factors were identified from MA07 using the VFAnalyser component of Virulence Factor DataBase (VFDB) [37]. Genes were visualised in MA07 using the JBrowse application on Galaxy Australia [38] and mapped to the genome map using SnapGene software. Further identifying information about predicted virulence factor sequences was determined by BLASTp search against non-redundant protein sequences (expect value 0.05). Where no similar sequences were found, protein structure was predicted from amino acid sequence using SWISS-MODEL, to identify proteins with similarity based on predicted tertiary structure [39]. To characterise the localisation of the identified MA07 VFs within the genome, online software Proksee (proksee.ca) was used to generate a circular genome map of MA07 and display the sequence similarity of MA07 to *M. abscessus* ATCC19977 and *M. tuberculosis* H37Rv.

### DNA extraction, library preparation and long-read sequencing

DNA extraction was performed as previously described using the DNeasy UltraClean Microbial kit (Qiagen), with some modifications [40]. After growth of MA07 on 7H10 agar, cells were vortexed with powerbeads for 5 minutes to increase bacterial lysis and improve DNA yield. Following DNA extraction, quality was assessed using Nano-300 spectrophotometer (AllSheng, China) and DNA concentration determined with a Qubit 2.0 Fluorimeter using the dsRNA BR Assay Kit (ThermoFisher). Following quality control, library preparation for long read sequencing was performed using the Rapid Barcoding Kit (SQK-RBK004) and sequencing was performed on the GridION platform on an R9.4.1 flow cell (Oxford Nanopore Technologies plc, UK). Base-calling was performed using Guppy version 3.4.5 + f11b1b. Hybrid assembly was performed with Unicycler v0.4.8 [41] and annotation performed with Prokka v1.14.6.

### DNA and RNA extraction and qRT-PCR analysis

For genomic DNA extraction of *M. abscessus*, bacteria were grown to log phase in complete 7H9 and pelleted by centrifugation. Pellets were resuspended in solution 1 [25% (w/v) sucrose, 50 mM Tris), 50 mM EDTA + Lysozyme 500 μg/mL (Sigma)] then incubated at 37°C shaking (200 RPM) overnight. One volume of freshly prepared solution 2 [100 mM Tris, 50 mM EDTA, 1% sodium dodecyl sulfate (SDS; Sigma)] was added and incubated for four hours at 55°C, followed by addition of sodium chloride to a final concentration of 0.83 M. DNA was extracted by two rounds of mixture with phenol/chloroform/isoamyl alcohol extraction followed by precipitation of nucleic acids at – 20°C overnight using ice cold ethanol. Concentration and purity of DNA was determined by using Nano-300 spectrophotometer (Allheng, Hangzhou, China).

RNA extraction was performed using Qiagen RNeasy Mini Kit as per manufacturer’s instructions. For qRT-PCR analysis, genomic DNA was removed using TURBO DNA-free kit per manufacturer’s instructions followed by cDNA synthesis using tetro cDNA synthesis kit (Meridian bioscience, USA). After validation of RNA purity by PCR, qRT-PCR was performed using the sensifast-SYBR green NO-ROX kit and primers described in Table S4. Analysis was performed on the ROCHE Lightcycler 480 II at the University of Sydney, Australia using the following conditions: 95°C for 10 minutes, 50 cycles of 95°C for 10 s and 72°C for 5 s. Fold change in gene expression was normalised to the reference gene SigA (Laencina et al., 2018) and determined by calculating 2^(-ΔΔCT)^ or 2^(-ΔCT)^.

### Genome assembly, annotation and visualisation

FASTQ files generated from short-read sequencing were uploaded to the Galaxy Australia platform for quality control, assembly and annotation (usegalaxy.org.au; Afgan et al. 2018). Genome assembly was performed using SPAdes [43] and annotation performed using Prokka [44]. Contigs were aligned to the *M. abscessus* FLAC013 reference genome using progressive Mauve to create a singular FASTA file for BLAST analysis [45]. Short-read sequencing data were used for initial VF screening using VFDB . Circular genome maps showing similarity between different plasmids were created using Blast Ring Image Generator (BRIG v0.95;) using pMA07 as the reference sequence [47].

### Phylogenetic analysis

Contigs and raw sequence read data for 1,160 isolates (sourced from Genbank or generated by Illumina sequencing by CIDM) were mapped against the plasmid using Snippy (version 4.4.5) (https://github.com/tseemann/snippy). The snippy-core command was used to generate an alignment of core-genome single nucleotide polymorphisms (SNP) that formed the input for FastTree (version 2.1.10) to generate a maximum likelihood tree using the general time reversible model. [48] Read-mapping coverage across plasmid genes was assessed for each isolate using bedtools (v2.30.0) to determine the presence of plasmid genes.[49]. The read coverage for each gene across the isolates analysed was determined using NumPy version 1.19.2 [50] and pandas version 1.1.5 [51]. The newick tree and read coverage of each strain for the plasmid was visualised using interactive tree of life (iTOL) version 6 [52]. Accession numbers of sequences included in analysis are included in supplemental files.

### CRISPRi inducible knockdown

CRISPRi plasmids were generated by the digestion of 2 μg PLJR962 plasmid backbone [33] sourced from Addgene with EcoRI-HF and SAlI-HF followed by ligation of 200 ng digested vector with 2-4 ng sgRNA using sgRNA guide sequences described (Table S5). Plasmids were transformed into DH5α *E. coli* using the heat shock method plated on LB agar containing 50 μg/mL kanamycin. Constructed plasmids were purified using the Nucleobond Xtra Midi kit (Macharey Nagel, Düren Germany) then verified by Sanger sequencing and fragment analysis at the Garvan Institute of Medical Research, Sydney, Australia.

For transformation of *M. abscessus*, bacteria at a log phase of growth were washed with sequentially decreasing volumes of wash buffer (sterile de-ionised water with 10% glycerol and 0.02% tyloxapol), then electroporated with 0.1-5 μg relevant plasmid using two rounds of pulsation (2.5 kV) with a micropulser (Bio-Rad laboratories). Transformants were incubated for 3-4 hours while rolling in 7H9 at 37°C before plating on 7H10 agar with 50 μg/mL kanamycin. Plates were checked for the presence of colonies after 5-6 days, and colonies screened for successful incorporation of the plasmid by PCR and agarose gel electrophoresis, using primers PLJR962-F1 and PLJR962-R1

### Cell infection experiments

RAW 264.7 cells were seeded at a density of 5 X 10^4^ -1 X 10^5^ cells/mL in 96 well plates and allowed to adhere overnight, while THP-1 cells were differentiated for 72h in the presence of 100 ng/mL PMA. Cells were infected with *M. abscessus* at an MOI 2-10 in the presence of 500 ng/mL anhydrotetracycline (ATc) where required. After 3 hours, cells were washed with PBS and fresh complete media containing 250 μg/mL amikacin (Sigma) to limit *M. abscessus* extracellular growth. Cells were incubated for a further 1 hour, then washed with PBS and lysed with sterile TDW for CFU enumeration. Cell lysates were serially diluted and plated on LB agar to enumerate intracellular bacterial burden. For analysis of ESX components in THP-1 macrophages by qRT-PCR, 3×10^6^ THP-1 macrophages per replicate were infected with *M. abscessus* MA07 at an MOI 10. At the indicated time points, cells were washed with PBS followed by lysis with RNAse free H_2_O and pelleting of bacteria by centrifugation (2000xg, 30 minutes, 4 °C) and storage in Trizol at -80°C for later analysis.

## Acknowledgements

The authors would like to thank Professor Nick West for helpful guidance in using the CRISPRi knockdown strategy.

## Competing interests

The authors declare no competing interests.

## Funding

This work was supported by the NHMRC Centre of Research Excellence in Tuberculosis Control (APP1153493).

## Authors contributions

KCF, CC and JAT conceived and planned the experiments; KCF, AHB, SW, SA, AB, ES and EM performed experiments and analysis; KCF, AHB, VS, CC, TPS and JAT contributed to interpretation of the results; KCF drafted the manuscript with contributions from JAT, AHB and TJS; all authors contributed to the final manuscript.

## Data availability

The datasets used and/or analysed during the current study are available from the corresponding author on reasonable request.

## Notes

### Competing Interest Statement

The authors have declared no competing interest.

